# Signals from the brood modulate the sleep of brood tending bumblebee workers

**DOI:** 10.1101/500744

**Authors:** Moshe Nagari, Ariel Gera, Sara Jonsson, Guy Bloch

## Abstract

Sleep is ubiquitous in vertebrates and invertebrates, and its chronic lost is typically associated with reduced performance, health, or survival. Nevertheless, some animals can give up sleep in order to increase survival or mating opportunities. We studied the interplay between sleep and brood care in the social bumblebee *Bombus terrestris*. We first used video recording and detailed behavioral analyses to confirm that the bumblebee shows the essential behavioral characteristics of sleep. Based on these analyses we next used immobility bouts of >5′ as proxy for sleep in automatic activity monitoring records, and found that sleep is severely reduced in the presence of larvae that require feeding or pupae that are not fed. Reduced sleep was correlated with wax pot building, which is a behavior typical to nest founding mother queens. Sleep was also reduced in the presence of empty cocoons, but this effect was transient and reduced with time. This observation that is consistent with the presence of a sleep modulating pheromonal signal. These results provide the first evidence for brood modulation of sleep in an insect, and are consistent with the hypothesis that plasticity in sleep can evolve as a mechanism to improve care for dependent juveniles.

## Introduction

Sleep is an arousal state ubiquitous in vertebrate and invertebrate animals. The sleep state is characterized by extended bouts of behavioral quiescence with a distinct body posture and reduced responsiveness to external stimuli (reviewed in Cirelli and Tononi, 2008; see also Nath *et al.*, 2017; Omond *et al.*, 2017; Raizen *et al.*, 2008; Vorster *et al.*, 2014). Sleep is regulated by the circadian clock, that controls the daily timing of sleep, and by homeostatic mechanisms; extended periods of arousal increase the drive for sleep and can increase subsequent sleep duration or intensity (Allada et al. 2017; Blum et al. 2018; i.e., slow-wave activity during sleep). The ubiquity of sleep and the evidence for homeostatic need to compensate for lost sleep suggest that it is functionally important. Indeed, in many animals it was shown that extended periods of sleep deprivation or sleep fragmentation led to impaired cognitive performance or health (Beyaert et al. 2012; Dawson & Reid 1997; Hussaini et al. 2009; Klein et al. 2010; Rolls et al. 2011; Vorster & Born 2015; Vyazovskiy et al. 2011). In spite of this evidence, the functional significance of sleep as well as its evolutionary origin remain debated issues.

One set of evidence adding to the mystery surrounding the adaptive function of sleep is that many animals can naturally show extended periods of activity with little or no sleep (Eban-Rothschild et al. 2017). For example, when foraging for food (Rattenborg et al. 2016), under predation risk (Dominguez 2003; Lendrem 1984; Lesku et al. 2008; Rattenborg et al. 1999), during seasonal migration (Fuchs et al. 2009; Jones et al. 2010; Rattenborg et al. 2004), and during the breeding season (Lesku et al. 2012). Activity around the clock with attenuated circadian rhythms and reduced sleep has been also reported for animals tending offspring (Bloch et al. 2013). For example, Killer-whale and bottle-nose dolphin mothers show attenuated circadian activity rhythms and little sleep during the first month post-partum, when they need to help their calves surface to breath (Lyamin et al. 2005). Similarly, mother rats show decreased non-REM sleep duration on the days following parturition when their newborn pups are completely dependent on their nursing and care (Sivadas et al. 2017). Interestingly, the mother’s sleep is deeper when she eventually falls asleep, possibly compensating for the reduction in sleep duration. Actigraphic studies on human mothers during the first few weeks post-partum suggest a similar attenuation in circadian rhythmicity and increased wake activity after sleep onset (i.e., interrupted night sleep) compared to the pre-partum period (Nishihara et al. 2002; Nishihara & Horiuchi 1998). The activity of the mother was correlated with the baby movements during the night, suggesting that sleep loss was linked to maternal care (Nishihara & Horiuchi 1998). Taken together, these studies suggest profound plasticity in sleep and circadian rhythms in animals that progressively care for their young offspring.

In the social hymenoptera (bees, ants, and wasps), very similar association between reduced sleep duration, attenuated circadian rhythms, and care for the young, is shown by sterile workers tending brood in their colony. Social insects live in colonies and demonstrate division of labor between the workers, who provide all colony needs. The best studied division is between foragers who collect resources outside the nest, and nurses, who maintain the nest and tend the brood. Brood care behavior is associated with attenuated circadian rhythms, whereas foragers have strong circadian rhythms that are needed for time compensated sun-compass orientation and timing visits to temporally restricted food sources such as flowers (Eban-Rothschild & Bloch 2012). In honeybees, it was shown that this task-related plasticity in circadian rhythms is socially regulated. Nurse bees that come in close contact with both larvae, which they feed, and pupae, which do not require feeding, show around the clock activity and attenuated circadian rhythms (Nagari et al. 2017; Shemesh et al. 2010). These observations are consistent with the premise that around-the-clock activity improves brood care, for example, by enabling recurrent feeding of larvae, or improved thermoregulation. However, such frequent brood tending may come with a cost of reduced sleep.

Given that sleep is circadianly regulated and that nurse bees care for brood around the clock, it can be hypothesized that sleep is reduced in brood tending bees. The sleep (or “sleep-like”) behavior of the honeybee has been characterized independently by several groups; collectively they suggest that honeybees show a typical sleep behavior with reduced muscle tonus and elevated arousal threshold (Eban-Rothschild & Bloch 2008; Kaiser 1988; Sauer et al. 2003; Kaiser & Steiner-Kaiser 1983). A few studies explicitly focused on sleep like behavior in bees that are active around-the-clock (Eban-Rothschild & Bloch 2008; Klein et al. 2008; Klein et al. 2014). These studies suggest that around the clock active honeybees do not sleep less than foragers do, but rather their sleep is more intermitted. We previously examined the influence of pupae on the locomotor activity of isolated nurses in the laboratory using automated activity monitoring and found that overall sleep amount was not affected by the presence of brood, despite an effect on circadian rhythms (Nagari et al. 2017). These findings suggest that although the brood attenuates circadian rhythms in nurse honeybees, their overall daily sleep amount is not affected.

The relationship between brood care and circadian rhythmicity has been previously explored in the incipiently eusocial bumblebee (*Bombus terrestris*) (Eban-Rothschild et al. 2011; Yerushalmi et al. 2006). Unlike honeybee queens, who lay eggs but do not care for the brood, bumblebee queens found their nests alone and thus need to provide all the needs of the first batch of brood. Nest-founding queens show attenuated circadian rhythms in the presence of brood, but rapidly switch to activity with circadian rhythms when the brood is removed (Eban-Rothschild et al. 2011). Moreover, broodless queens that lay again after the brood removal switch to activity with attenuated circadian rhythms. Remarkably, this switch occurs even before the eggs are laid, suggesting that both the brood presence and the reproductive physiology of the queen modulate circadian rhythmicity. The emergence of the first batch of worker bees mark the beginning of the social phase of the colony cycle during which, the workers take on the brood tending duties and the queen focuses on egg-laying (Woodard et al. 2013). Similarly to honeybee nurses, and to nest founding bumblebee queens, nurse bumblebee workers tend the brood around the clock (Yerushalmi et al. 2006). It is not known, however, whether they also sleep less.

Here, we asked whether brood at different developmental stages affects sleep duration in bumblebee workers. We also asked whether the worker sleep is influenced by their reproductive physiology. First, we performed detailed video recordings to characterize the sleep-like behavior of individually isolated *B. terrestris* workers. Based on these analyses we defined a period of 5 minutes of immobility as a proxy for sleep. We next used this proxy to monitor sleep with an automated video-based monitoring system and test the following hypotheses: (1) Workers that care for brood have reduced daily sleep. (2) Larvae, which require feeding, have a stronger effect on the worker sleep compared to non-feeding pupae. (3) The brood effect on worker sleep is mediated by pheromones. (4) Activity around the clock and reduced sleep is associated with increased ovarian activity and wax building. We monitored individually isolated bumblebee workers in the presence of various brood stages or brood related stimuli.

## Results

### Experiment 1: Detailed analyses of the behavior of workers isolated with or without pupae

Given that motionless animals can be at various arousal states, we first used video recording and detailed analyses to characterize the behavior of immobile bumblebee workers. Bouts during which bees appeared immobile but were partially hidden were not included in these analyses (“Immobile unknown” category; see Table 1). Bees with a pupa (‘Pupa+’) spent significantly more time on the pupa compared to the time broodless bees (‘Brood-’) spent on the piece of wax (0.3 ± 0.1, n = 18 and 0.13 ± 0.06, n = 19, respectively, means ± s.e.m; Two-sample t-test, *p = 0.007*). In some cases, the bees were observed grooming, inspecting, or touching the pupa with their antennae and mouthparts. In other cases, they displayed a distinct behavior, during which they extended and continuously vibrated their abdomen in pumping movements, while clutching their abdomen to the pupa and wrapping their legs around it. This behavior was similar to the incubation behavior that was first described in detail by Heinrich, and accordingly we termed it “Incubation like” (Heinrich, 1972a, 1972b, 1974; see Movie 1 and Table 1). Incubating bees were typically immobile, but occasionally moved and changed their position. Their antennae were typically extended with an angle of > 90° between the frons and scape and they moved with a higher frequency compared to bees performing sleep-like behavior (Fig. 1A, two-sample t-test; Movies 1 and 2). During sleep-like bouts, the bees had a more relaxed body posture, their abdominal ventilation pumping was discontinuous, their antennae had an angle of < 90° between the frons and scape, and the antennal movements were less frequent (Fig. 1A, Movie 2).

**Figure 1.**
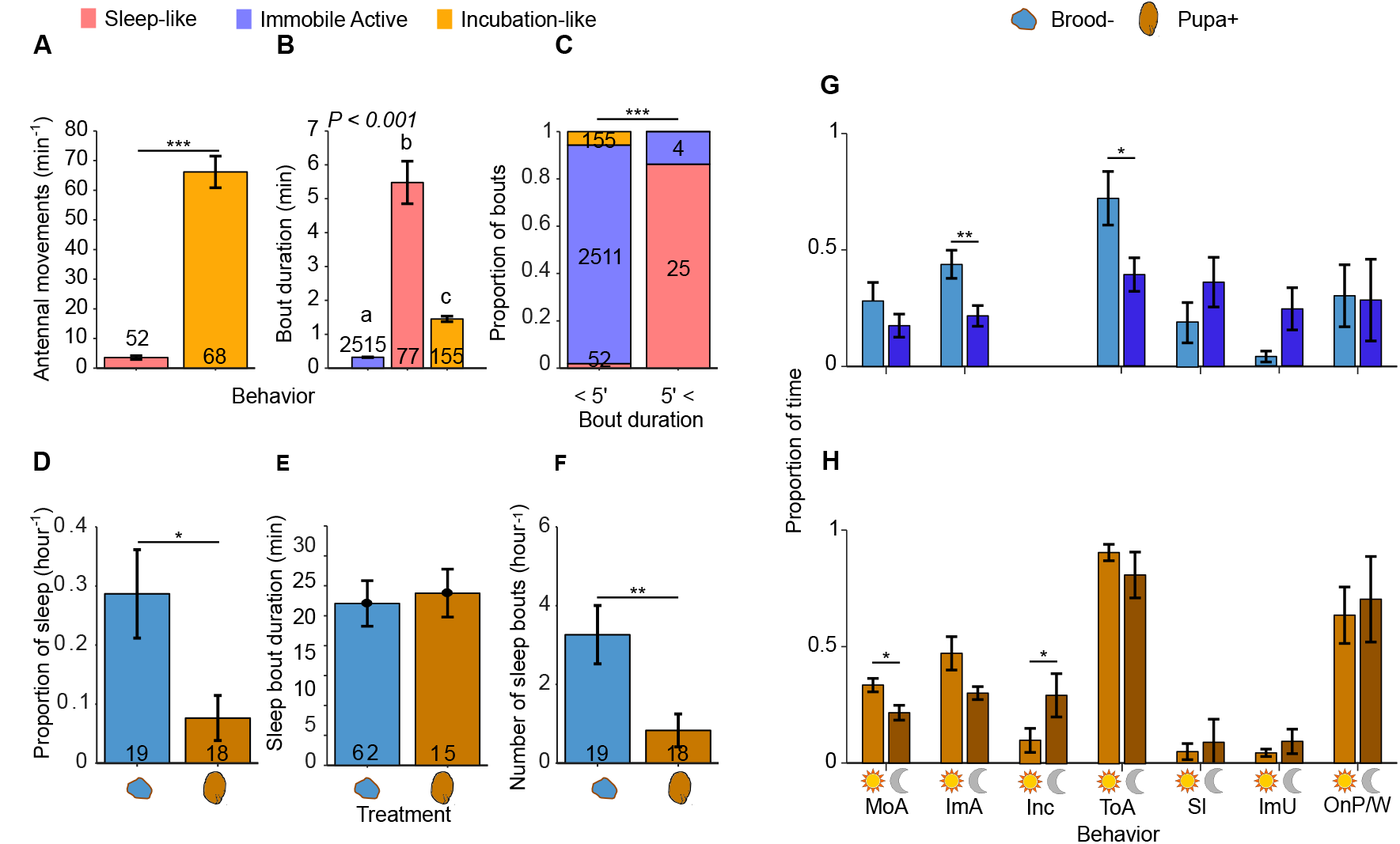
Video based behavioral analyses of bees isolated with or without a pupa. (A—C) Characterization of the behavior of immobile bees. (A) Antennal movements during bouts of ‘Sleep-like’ and ‘Incubation-like’ behaviors (means ± s.e.m, n; two-sample t-test). (B) Bout duration (means ± s.e.m, n; Kruskal-Wallis test followed by Mann-Whitney tests with Bonferroni corrections). (C) The proportion of bouts in which immobile bees showed different behaviors, for bouts lasting ≥ 5’ or < 5’ (X^2^ test for independence; sample sizes inside the bars). (D—F) The influence of pupae on the ‘Sleep-like’ behavior of worker bees. (D) Proportion of ‘sleep-like’ behavior during the video recordings. (E) Bout duration. (D) Number of sleep bouts. The plots show the means ± s.e.m, sample-sizes and the results of two-sample t-tests (D, F) or Mann-Whitney test (E). (G, H) The proportion of time each behavior was performed during the day and the night. (G) bees without pupa. (H) bees with a pupa (mean ± s.e.m; n = 5 bees for each treatment; paired t-test). Light and dark bars for each behavioral category indicate subjective day and night, respectively. MoA: Mobile-active; ImA: Immobile-active; Inc: Incubation-like; ToA: Total-activity; Sl: Sleep; ImU: Immobile-unknown; OnP/W: On-pupa/wax. *– *p* < *0.05*; **– *p* < *0.01*; ***– *p* < *0.001* in all plots.

**Table 1.** The behavioral Categories used for video analyses in experiment 3

| Behavior | Description |
| --- | --- |
| **On pupa/wax** | Stayed on the pupa or control wax stimulus for at-least 5 consecutive seconds |
| **Mobile active** | Changed position for $\geq 5$ consecutive seconds |
| **Immobile active\*** | Immobile (no change in position for $\geq 5$ consecutive seconds). Antennae raised at $> 90$ degrees between frons and scape. Antennae or head constantly moving. Abdomen continuously pumping |
| **Immobile unknown\*** | Immobile but behavior could not be determined because bee was hidden |
| **Sleep-like\*** | Immobile. Antennae are lowered at $\leq 90$ degrees between frons and scape. Antennae moving at a frequency of $< 20$/sec. Discontinuous abdominal pumping. Relaxed body posture. |
| **Incubation-like\*** | Immobile. Abdomen 'smeared' on pupa, vigorously pumping. Legs wrapped around the pupa. Antennae raised with $> 90$ degrees between frons and scape and constantly moving. Head and antennae not touching the substrate. |
| **Total activity** | All the non sleep activities combined (not including Immobile unknown). |

\* Behaviors with relative immobility

‘Immobile active’ bees were clearly awake, showing behaviors such as handling wax or tending the pupa, but they did not continuously change their position (see Table 1). The mean bout duration (Kruskal-Wallis test followed by Mann-Whitney *post-hoc* comparisons and Bonferroni corrections, Fig. 1B) and bout lengths distributions (Fig. S1A; Two-sample Kolmogorov-Smirnov test) significantly differed between sleeping, incubating, and awake immobile bees, indicating that these behaviors may be separated based on the bout length index. After visually examining bout lengths distributions (Fig. S1A, B), we tested whether a cutoff of 5’ bout duration can distinguish sleep bouts from immobile-active and incubation-like bouts in bees with or without a pupa. Our analyses revealed that 91.1% of the immobile bouts lasting more than 5’ were performed by bees in the sleep-like state (Fig. S1B; 100% and 88.7% for bees with or without a pupa, respectively). This corresponded to 86.6% of the number of > 5’ bouts (100% and 81%, respectively; Fig. 1C). Importantly, we did not observe any incubation-like bouts lasting more than 5’. One third (33.3%) of the total sleep-like time was consisted of bouts shorter than 5’ and thus may not be detected using the ≥ 5’ of immobility index (Fig. 1C; 24% and 36% for bees with or without a pupa, respectively). However, sleep-like bouts constituted only 1.9% of the total short immobile bouts (i.e., sleep-like, incubation-like and immobile-active bouts; 0.4% and 4.8% for bees with or without a pupa, respectively; Fig. 1C), suggesting that lowering the threshold below 5 minutes may significantly increase false detection of sleep-like bouts. Based on these analyses ≥ 5’ immobility bouts lasting more than 5’ constitute a conservative index for sleep that can be used to estimate sleep duration in data collected by automatic locomotor activity monitoring. However, given that our detailed video analyses suggest that about 33% of the sleep duration is performed in bouts shorter than 5’, for a more accurate estimation of total sleep duration it is necessary to multiply the total duration of immobility bouts > 5’ by 1.5. These results suggest that for bees without a pupa the > 5 ‘ index produces more false detection of sleep compared to bees with a pupa. However, during ‘Immobile active’ bouts, bees typically display discontinuous movements and we assume that most of these movements would be detected with our automated monitoring system that is based on continuous video recording (see Methods).

We next compared bees housed with or without a pupa and found that in the presence of pupa bees slept less. All 5 broodless bees and 3 of 5 of the bees with a pupa were observed sleeping in at least one recording session. The proportion of sessions with no sleep (14 of 18 and 6 of 19 and in bees with or without a pupa, respectively; X^2^ test, *p = 0.016*) and the overall proportion of time asleep during the video recording sessions (Fig. 1D; two-sample t-test, *p = 0.016*) were significantly reduced in bees with a pupa. In addition, the number of sleep bouts, but not the sleep bout duration, was reduced in bees with a pupa (Fig. 1D, E; two-sample t-test and Mann-Whitney test; *p = 0.049* and *p = 0.18*, respectively). A detailed comparison of the day and night observations further suggests that the temporal organization of behavior was affected by the presence of a pupa. In broodless bees, the proportion of Immobile-active and Total-activity (the combination of all wake activities, see Table 1), but not Mobile-active, behaviors was significantly increased during the subjective day compared to the subjective night (Fig, 1G; paired t-tests). The proportion of Sleep-like behavior, as well as immobile bouts in which we could not unambiguously assign a behavior (‘Immobile-unknown’; Fig. 1G; Table 1), appeared higher at night in broodless bees, but this trend did not cross the statistical significance threshold (*p = 0.34* and *p = 0.07*, respectively). Bees that were housed with a pupa were overall similarly active during the day and the night (‘Total-activity’ in Fig. 1H; *p = 0.35*). However, a finer examination revealed that they showed more ‘Mobile-active’ behavior during the day, and a similar trend for ‘Immobile-active” behavior (P=0.068; Fig. 1H). This apparent discrepancy may be explained by the finding that Incubation-like behavior was significantly higher during the night sessions (Fig. 1H). These video analyses show that the presence of pupae affects the temporal organization of behavior and the total amount of sleep.

### Experiment 2: The effect of larvae on the sleep of tending workers

Based on the detailed observations in Experiment 1, we automatically monitored the locomotor activity of isolated workers and used the > 5’ bout of inactivity as a proxy for sleep in all following experiments. We first tested the influence of both male larvae (offspring of a queenless worker) and female larvae (offspring of a mated queen) on the amount of sleep in individually isolated tending workers (see experimental design in Fig. 2A). Seven of the 35 control workers that were isolated with no brood (‘Brood-’) laid eggs during the monitoring session. Egg laying did not significantly affect the proportion of daily sleep (0.24 ± 0.066 and 0.34 ± 0.036, mean ± s.e.m in layers and non-layers, respectively; independent t-test, *p* = *0.23*). However, egg-laying broodless workers had weaker circadian rhythms in locomotor activity compared to broodless workers that did not lay eggs (power of circadian rhythms = 52.8 ± 26 and 144 ± 25.9, *p* = *0.022*). Given this trend and the possible interaction between circadian rhythmicity and sleep, we excluded the egg-laying workers from the following sleep analyses. Using the remaining data set, we found that the presence of larvae had a significant influence on the total amount of sleep, but there was no significant effect for the larvae gender or whether the focal be experienced queenright or queenless conditions before isolation and monitoring (Social experience factor; Fig. 2C, two-way ANOVA, *p-values* are shown in the figure).

**Figure 2.**
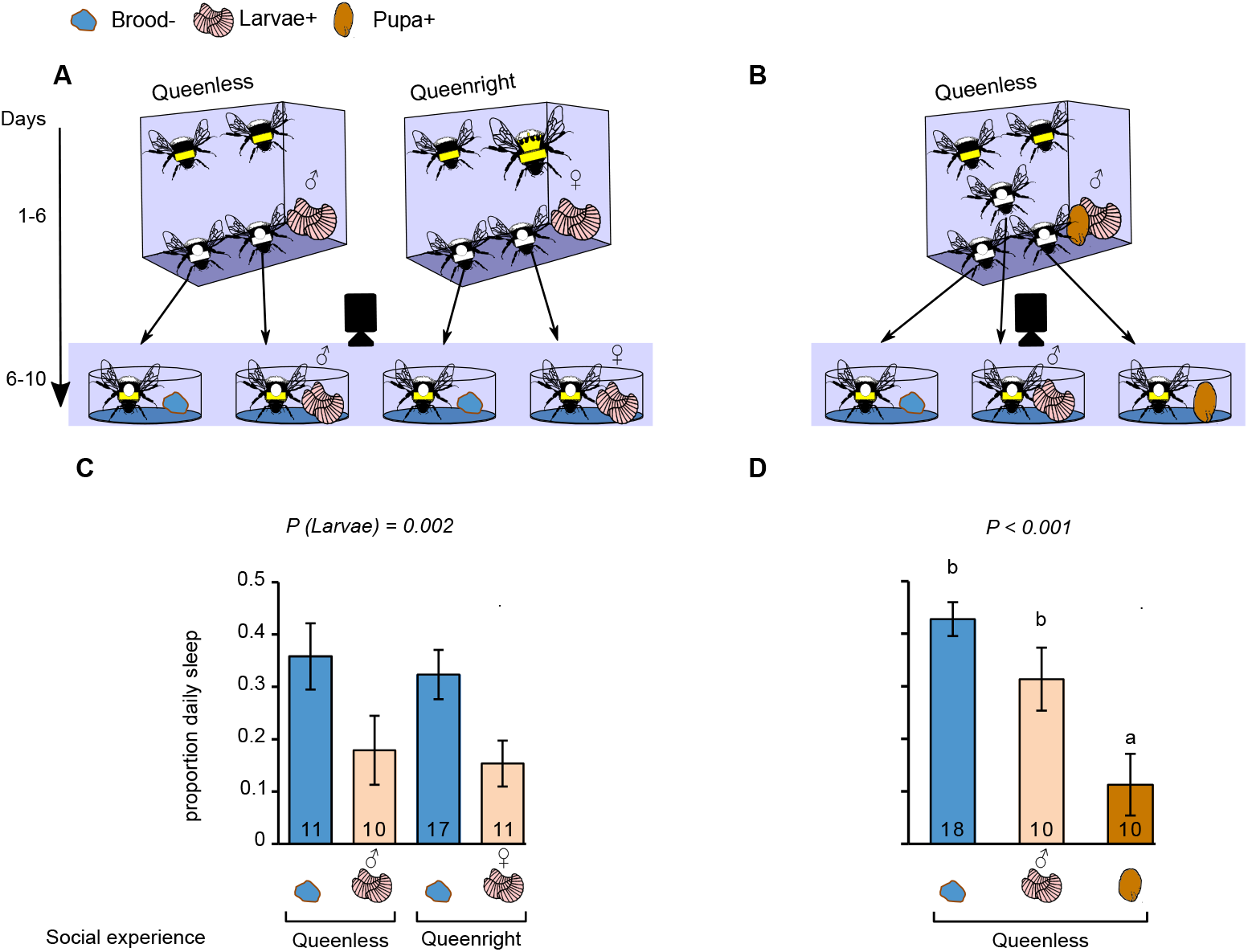
The effect of live brood on the sleep duration of tending workers. (A) The experimental setup of Experiment 2. Pairs of sister callow bees (indicated with white thorax color and a white dot) were introduced into queenless (top left) or queenright (top right) cages on Day 1 or 2 and kept under constant darkness and temperature. On Day 6 (bottom panels), each focal bee was individually isolated with either a piece of wax (‘Brood-’) or with ~8 male or female larvae (‘Larvae+’) from her queenless or queenright cage, respectively. (B) The experimental setup of Experiment 3. Trios of sister callow bees were introduced into queenless cages on Day 1 or 2. On Day 5, a pupa was introduced into each cage. On Day 6, each focal bee was individually isolated with a piece of wax (‘Brood-’), ~8 larvae or a single live pupa (‘Pupa+’) from her source cage. Timeline for (A) and (B) is shown to the left. (C) The proportion of daily sleep for bees in Experiments 2 (means ± s.e.m, n). Two-way ANOVA with the factors Larvae and Social experience in the cage before isolation; *p*(*Social experience*) = *0.58, p*(*Larvae X Social experience*) = *0.93*. (D) The proportion of daily sleep for bees in Experiments 3. One-way ANOVA; treatments marked with different letters are statistically different in a LSD *Post-Hoc* test.

### Experiment 3: The effects of larvae and pupae on the sleep of tending workers

Here we compared the effects of larvae to that of pupae, which do not require feeding, on the sleep of brood tending workers. Given that in Experiment 2 larvae sex and previous social experience did not affect the amount of sleep, we placed the focal bees with male larvae, which were the offspring of queenless workers (Fig. 2B). None of the focal bees in this experiment laid eggs during the monitoring session. The treatment affected the amount of sleep, with the pupa having a stronger effect compared to that of larvae (Fig. 2D; one-way ANOVA followed by LSD *Post-Hoc* test). The sleep duration in workers with larvae was not significantly lower compared to control bees housed with a piece of wax, although there was a clear trend in this direction (Fig. 2D). Given the strong influence of the pupa on sleep we decided to investigate the pupae effect with more detail.

### Experiment 4: The effects of live pupae and empty cocoons on the sleep of tending workers

To test whether a live pupa is required to affect the worker sleep, we compared the effects of an empty cocoon from which the pupa was removed (‘Empty’); a live intact pupa (‘Pupa+’); a sham-treated cocoon with a live pupa (‘Sham’) or bees without brood (‘Brood-’; Fig. 3A). Some of the bees laid eggs during the monitoring sessions (1, 2, 3, 3 and 4, 1, 0, 2 from the ‘Brood-’, ‘Empty’, ‘Sham’ and ‘Pupa+’ treatments in trials 2 and 3, respectively; none of the bees laid eggs in Trial 1). The bees that laid eggs showed a trend for reduced sleep, but this was not statistically significant in both trials (Trial 2: 0.078 ± 0.029 and 0.16 ± 0.031, two-sample t-test *p* = *0.063*, Trial 3: 0.084 ± 0.032 and 0.16 ± 0.028, *p* = *0.11*). Egg laying bees also showed significantly weaker circadian rhythms in locomotor activity in one of two trials (Trial 2: 61.6 ± 13.3 and 140.1 ± 30.4, *p* = *0.026;* Trial 3: 98.28 ± 30.29 and 196.89 ± 38.6, *p* = *0.058*). Given these putative effects, we excluded egg-layers from further analyses. In Trials 2 and 3, we recorded whether the bees built a wax pot during the experiment. We found that bees housed with a live pupa were significantly more likely to build a wax pot compared to bees housed without a pupa in Trial 3 (*χ*^2^ test for independence, *p* = *0.002;* Fig. 3B) and a similar nearly significant trend was observed in Trial 2 (*p* = *0.053*). Wax building behavior affected sleep: in a pooled sample of bees from all treatments, wax-pot builders tended to sleep less (two-sample t-test; insets in Fig. 3C).

**Figure 3.**
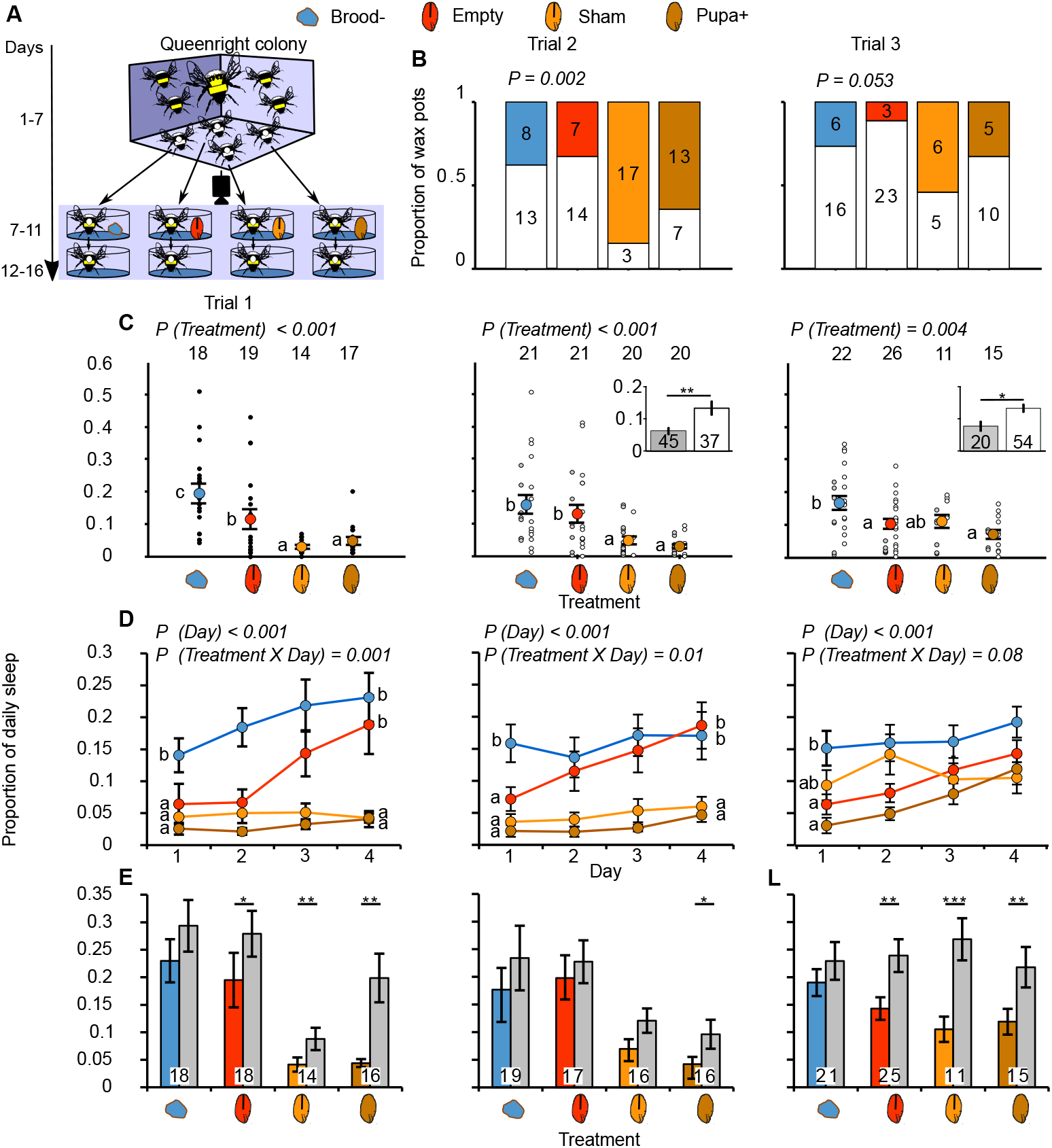
The effect of pupae and empty pupal cocoons on the sleep duration of tending workers. (A) Experimental setup for Experiment 4. Paint marked sister callow bees were introduced into queenright colonies on Day 1 or 2 (top panel). On Day 7, each bee was transferred into an individual cage with one of the following treatments (middle panel): a piece of wax (‘Brood-’), an empty cocoon from which the pupa was removed (‘Empty’) a sham treated cocoon with a live pupa (‘Sham’), or a live intact pupa (Pupa+). After five days of monitoring, the treatment stimuli were removed and activity was monitored for 5 additional days (bottom panel). (B) The proportion of wax pot building on the 2^nd^ and 3^rd^ trial (not monitored in Trial 1); the *p*-values summarize X^2^ tests of independence. (C) The proportion of daily sleep on Days 8—11.black filled small circles in Trial 1 show values for individual bees; open and grey filled small circles In Trials 2 and 3 mark bees that did or did not initiate wax-pot building, respectively; color-filled circles with bars show means ± s.e.m, sample sizes are shown in the top. The letters indicate the results of repeated measures ANOVA (see below) followed by LSD *Post-Hoc* tests for the Treatment effect. The insets summarize the proportion of sleep for bees that built (grey) or did not build (open bars) wax pots across all treatments (means ± s.e.m; two-sample t-test). (D) The proportion of daily sleep across time (means ± s.e.m). The small letters indicate the results of One-way ANOVA followed by LSD *Post-Hoc* tests comparing the proportion of daily sleep on the 1^st^ and 4^th^ days of monitoring (Days 8 and 11 of the experiment). The *p-* values in C and D were obtained from two-way repeated measures ANOVA with the factors Treatment and Day. (E) The proportion of daily sleep (means ± s.e.m, n) on the day before (colored bars) and after (grey bars) removing the treatment stimuli (means ± s.e.m, n; paired t-tests). *– *p* < *0.05*; **– *p* < *0.01*; ***– *p* < *0.001* in all plots.

We found that sleep duration was affected by treatment in all three trials (Fig. 3C—E). Bees housed with a live pupa (‘Sham’ and ‘Pupa+’) slept less than the ‘Brood-’ bees that were housed with only a piece of wax (Fig. 3C, two-way repeated-measures ANOVA followed by LSD *Post-Hoc* tests; in Trial 3, the ‘Sham’ treatment effect did not reach the statistical significance threshold). The effect of empty cocoons was intermediate; bees subjected to this treatment slept less than bees housed with a piece of wax in Trials 1 and 3, with a similar but statistically nonsignificant trend in Trial 2. Additionally, these bees slept more than ‘Sham’ and ‘Pupa+’ bees in Trials 1 and 2. To better understand the intermediate effect of he empty cocoons, we examined the amounts of sleep on each day separately (Fig. 3D). We found that both ‘Day’ and ‘Day X Treatment’ interaction affected sleep (two-way repeated measures ANOVA; *p-values* are shown in the figure; in Trial 3, the interaction effect was *p* = *0.08*). The interaction effect stemmed mostly from a distinct pattern shown by bees housed with an empty cocoon: in the 1 day they slept significantly less than the ‘Brood-’ bees, and similarly to bees housed with a live pupa (both ‘Sham’ and ‘Pupa+’ bees; Fig. 3D; one-way ANOVA, *p = 0.01, p < 0.001* and *p* = *0.001* on Trials 1—3, respectively). Thereafter, their sleep amount gradually increased up to an amount comparable to ‘Brood-’ bees on the 4^th^ day. On Trials 1 and 2, bees with an empty cocoon slept significantly more on Day 4 compared to bees housed with alive pupa (*p* < *0.001, p* = *0.002* and *p* = *0.065* for Trials 1 —3, respectively). Thus, the empty cocoons had a sleep reducing effect on the first day, which lost its potency over time. Removal of the empty cocoon, sham treated pupa, and intact pupa treatments, but not the piece of wax (control treatment) led to a significant increase in sleep duration on the following day (paired t-tests, Fig. 3E; *p-values* are indicated in the figure. In Trial 2, the differences were statistically significant only for the intact pupa treatment). These results suggest that the cocoons release signals that affect sleep and lose their efficacy over time.

### Experiment 5: The effect of pupal extracts on the sleep of tending workers

The transient effect of the empty cocoons on sleep in Experiment 4 is consistent with the notion that it is mediated by pupal chemical cues that are absorbed to the cocoon and gradually lose their potency. We tested this hypothesis by exposing bumblebee workers to pupal extracts (‘Extract+’) or solvent alone (Vehicle’ treatment; See Fig. S2A) that were applied to washed cocoons. Intact pupae (‘Pupa+’) and pieces of wax (‘Brood-’) served as positive and negative controls, respectively. We repeated this experiment three times with different batches of focal bees. Two bees that laid eggs during the monitoring session on Trial 3 were excluded from the analyses (one each from the ‘Vehicle’ and ‘Pupa+’ treatments). As in Experiment 4, we also monitored wax pot building and found that in two of three trials bees with live intact pupae were more likely to build wax pots compared to the pooled negative control treatments (‘Brood-’ and ‘Vehicle’; Fig. S2B; *χ*^2^ test for independence followed by *Post-Hoc* comparisons). In Trials 2 and 3, bees that were housed with extract treated cocoons showed a similar, but weaker, trend that was statistically significant only in Trial 2 (Fig. S2B). In contrast, in Trial 1, the bees housed with only a piece of wax (‘Brood-’) were more likely to build wax pots compared to bees exposed to cocoon extracts (‘Extract+’ bees; Fig. S2B). In a pooled sample of bees from all treatments, we found that wax-pot builders slept less than bees that did not (Fig. S2C insets; two-sample t-test). In Trials 2 and 3, as in the other experiments, bees that were placed with a live pupa slept less compared to bees without a live pupa (Fig. S2C). The cocoon extracts, however, did not affect sleep duration in any of the trials. Thus, although the pupal extracts retained some weak effect on wax-pot building, at least in the 2^nd^ trial, they failed to mimic the effects of live pupae on the amount of sleep. This experiment further strengthens the association between wax building and sleep duration.

### The relationship between ovarian development and sleep

Given that egg-laying workers tended to have reduced sleep duration and weaker circadian rhythms, we tested the relationship between the presence of brood, ovarian state and sleep duration in a random sample of bees from Experiment 3 (10 ‘Brood-’, 7 ‘Larvae+’ and 4 ‘Pupa+’ bees) and in the bees from Trials 2 and 3 of Experiment 4. The ovaries of bees housed with live brood (larvae or a pupa; largest ovarian oocyte length mean ± s.e.m = 1.5 ± 1.2mm), were more developed compared to the ovaries of control bees housed with a piece of wax (1.12 ± 1.27mm; Fig. S3; two-way ANOVA, *p*(*Brood*) = 0.001, *p*(*Experiment*) = 0.006 and *p*(*Brood X Experiment*) = *0.91*) but ovarian state did not significantly affect sleep duration (multiple linear regression; *p*(*Oocyte length*) = 0.157 and *p*(*Experiment*) < *0.001*; Fig. S3). These analyses suggest that the presence of brood has a weak effect on the worker ovarian state, but this does not account for the observed effects on the worker sleep duration.

## Discussion

We characterized for the first time a sleep-like state in a bumblebee and showed that bumblebees that do not move for > 5’ are significantly more likely to be asleep rather than immobile and awake. Based on these findings, we used a > 5’ of immobility index to estimate sleep amount using high-throughput automated locomotor activity monitoring and found that sleep in bumblebees is strongly regulated by the presence of brood. Workers slept less when housed with larvae that require feeding as well as in the presence of non-feeding pupae. The presence of a pupa also resulted in an increased propensity to build wax pots and increased ovarian activation. Wax-pot building, but not ovarian state, was correlated with reduced sleep. Empty cocoons, from which the pupae were removed, induced a transient reduction in daily sleep suggesting that substances that are deposited onto the cocoons and lose their effect over time mediate the pupae effect on sleep. Taken together, these results suggest that the brood tending behavior of sterile workers is associated with their reproductive physiology, consistently with the reproductive groundplan hypothesis for division of labor. Importantly, the findings present the first evidence in an insect that the presence of brood modulates sleep and suggest that this effect is mediated by chemical signals. Additional efforts are needed for identifying the specific brood signals that mediate its effect on the worker sleep.

The sleep-like behavior of bumblebee workers shows the essential behavioral characteristics of sleep in other animals, and is particularly similar to that of the honeybee (Eban-Rothschild & Bloch 2008; Kaiser 1988;Kaiser & Steiner-Kaiser 1983 Klein et al. 2008). These similarities include bouts of immobility with a specific body posture and reduced muscle tonus. As in honeybees, the bumblebee sleep-like state is characterized by an angle of < 90° between antennal scape and frons, reduced antennal movements and discontinuous abdominal pumping (Movie 2). Sleep-like bouts constituted more than 85% of the immobile bouts longer than 5 minutes. This allowed us to use the duration of immobility bouts as a proxy for sleep. Our 5’ of immobility proxy for sleep is similar to that developed for honeybees and *Drosophila* (Eban-Rothschild & Bloch 2008; Shaw et al. 2000).

We show that the sleep-like behavior of bumblebee workers is strongly regulated by the presence of brood. The effect of pupae was confirmed both by direct observations using video recordings and by automated locomotor activity monitoring. The significant increase in the amount of sleep on the day after removing the pupae from the monitoring cages in Experiment 4 gives a strong causal support for the brood effect on worker sleep (Fig. 3E). There are reports that sleep is similarly reduced in the presence of juveniles in postpartum mothers of humans, rats and some cetaceans (da Rocha & Hoshino 2009; Lyamin et al. 2005; Nishihara & Horiuchi 1998; Shinkoda et al. 2002; Sivadas et al. 2017). For example, in humans, the presence of the baby in close proximity to the mother causes shorter and more interrupted mother sleep (McKenna et al. 1993; Nishihara & Horiuchi 1998; Volkovich et al. 2018). Similarly, in rat mothers, search of the pups for the mother’s nipples interrupts her sleep (da Rocha & Hoshino 2009); after weaning, the mother’s sleep returns to normal (Sivadas et al. 2017). Given that in humans the newborns are among the most neurologically immature at birth, the sleep loss of the mother is thought to be adaptive in allowing improved monitoring and care for the young (McKenna et al. 1993). Bumblebee brood is also completely dependent on its caregivers for survival but, in contrast with mammals, the tending bumblebee adults are not the mothers of the juveniles, but rather their older siblings.

One of the key findings of this study is that even the presence of empty pupal cocoons was sufficient to induce reduced sleep in tending workers. In all the three trials of Experiment 4, the effect of the emptied cocoons was similar to that of the live pupae on the first day of monitoring. However, the effect of the empty cocoons on worker sleep was transient, and after four days, the tending bees showed an amount of sleep comparable to that of bees with no brood. The most plausible explanation for this effect is that chemical substances from the pupae were left on the empty cocoons and modulated the nurses’ sleep. These chemicals then gradually degraded or evaporated over time. To further test this hypothesis, we exposed workers to washed cocoons impregnated with n-Pentane pupal extracts, which failed to mimic the pupal effect on sleep. However, these results do not reject the hypothesis that brood pheromones affect sleep. It is possible that some active components were not effectively extracted by the protocol we used, that these components lost their potency, or that the pheromonal effect depends on the presence of other chemicals (Funaro et al. 2018). This premise is supported by the finding that the pupal extracts retained only a partial effect on the propensity to build wax pots. Thus, future studies need to test additional extraction and application protocols. To our knowledge, these findings present the first evidence for brood pheromones affecting sleep.

The strong effect of the pupae indicates that brood feeding is not the only factor driving the reduction of sleep in workers because bumblebee pupae do not require feeding. Our detailed observations revealed that the pupae induced incubation-like behavior in isolated workers (Movie 1). Although we did not directly monitor heat production, the behavior that we observed was remarkably similar to Heinrich’s descriptions of incubating bumblebee (*Bombus vosnesenskii*) queens and workers (Heinrich 1972a; Heinrich 1972b; Heinrich 1974). He reported that the temperature of a nest foundress queen’s brood clump was tightly regulated during the night, but during the day the queen frequently left the brood and its temperature dropped (Heinrich 1974). Interestingly, consistent with these observations, we found that under constant ambient conditions, workers incubated brood more frequently at night, and moved more during the day (Fig. 1H). These findings suggest that brood incubation contributes to the reduction in sleep in tending workers, specifically to the more prominent sleep reduction at night (Fig. 1H). They also suggest that incubation behavior is regulated by an endogenous circadian clock. In honeybees, who also incubate their brood, the temperature of pupae is more tightly regulated compared with larvae (Kronenberg & Heller 1982). Pupae thermoregulation appears to be functionally significant in honeybees because even a slight deviation from optimal temperature impairs pupal development and the performance of the emerging adults (Groh et al. 2004; Tautz et al. 2003). In bumblebee colonies, the brood temperature is typically kept at 28—32°C throughout the day (Crall et al. 2018; Vogt 1986). The strong effect of pupae on sleep, together with the indications that incubation is increased at night in bumblebees at the expanse of sleep, suggest that brood incubation is one of the factors that drive nurses to reduce their sleep. Nevertheless, the bees showed additional brood tending behaviors such as construction of a wax pot next to the pupa and pupal grooming. Thus, it is likely that brood incubation is not the only activity associated to reduced sleep in brood tending bees.

We found that workers with pupae were more likely to build wax pots compared with workers that were housed with a control piece of wax. Moreover, workers that built wax pots slept less compared to those that did not (Fig. 3C). We also found that bees that built eggcups and laid eggs tended to sleep less and showed weaker circadian activity rhythms, although the effect on sleep was not statistically significant. Interestingly, *B. terrestris* queens show a similar association between egg-laying, nest construction and reduced circadian activity rhythms (Eban-Rothschild et al. 2011). Queens that laid eggs switched to around the clock activity before they started building eggcups and laid their second batch of eggs. Consistently, Heinrich described a “broody” state in *B. vosnesenskii* queens, in which nest foundresses increase their body temperatures during both day and night before they begin to build their nest and start laying eggs (Heinrich 1972a). These similarity between the physiology and behavior of nursing workers and mother queens is consistent the premise that the nurse behavioral state evolved from a maternal state in the solitary ancestor of bumblebees, consistent with the reproductive ground-plan hypothesis for the evolution of division of labor (Amdam & Page Jr. 2010; Eban-Rothschild et al. 2011; West-Eberhard 1987). Nevertheless, although workers with brood had more developed ovaries compared to control broodless bees, ovarian state was not correlated with the amount of sleep. This is consistent with the finding that removal of the ovaries did not affect the strength of circadian rhythms in bumblebee queen (Eban-Rothschild et al. 2011). Taken together, our results suggest that sleep loss is a component of maternal physiology that is expressed in bumblebee brood tending workers.

Our automatic activity monitoring and detailed video records indicate that brood tending bumblebee workers show very little amounts of daily sleep. Given the numerous evidence for the importance of sleep for function and survival (e.g., Rechtschaffen *et al.*, 1983; Shaw *et al.*, 2002), such little sleep can be expected to come with a cost. However, studies indicate that under some ecological conditions animals can show extended periods of very little sleep. For example, male pectoral sandpipers show extremely little sleep during their short mating season in the Arctic summer, which may help them in the competition for mating opportunities (Lesku et al. 2012). They show negative correlation between sleep duration and intensity, suggesting that they partially compensate for the loss of sleep-time. Some passerine birds sleep very little during the migration season in which they fly at night and forage for food during the day (Fuchs et al. 2009; Jones et al. 2010; Rattenborg et al. 2004). It was suggested that they can recover some of the sleep loss during periods of ‘drowsiness’, an arousal state during which their eyes are mostly open and their head moves, but their vigilance is reduced (Rattenborg et al. 2004). However, this may not fully compensate for sleep-loss during migration because there is evidence for impaired cognitive performance in migrating compared to non-migrating conspecifics (Jones et al. 2010). Great frigate birds were suggested to partially compensate for extreme sleep loss during long migration flights with sleep rebound when they get back to shore (Rattenborg et al. 2016). We did not find evidence for sleep rebound after removing the pupae in Experiment 4 (Fig. 3E), and it is yet unknown whether the reduced sleep of nursing workers is associated with compromised performance or health.

In sum, our study provides a remarkable example for ecologically relevant modulation of sleep. The sleep-like behavior of bumblebee workers is severely reduced in the presence of brood, including during stages that the brood does not need to be fed. The effect of brood is transient, and in the case of pupae appears to be at least partially mediated by substances deposited onto the cocoon and lose their efficacy over time. Brood tending workers show many similarities with mother queens.

## Materials and Methods

### Bees

*Bombus terrestris* colonies were obtained from Yad-Mordechai Pollination Services, Yad-Mordechai, Israel. Colonies were housed in wooden nest-boxes (30 × 23 × 20 cm) with transparent plastic covers. The bees were fed ad libitum with pollen (collected by honeybees) mixed with commercial sucrose syrup (Yad-Mordechai Pollination Services, Israel). The colonies were kept in an environmental chamber with constant temperature and humidity (27—29°C; relative humidity = 40—60%) and under constant dim red light (Edison Federal EFEE 1AE1 Deep Red LEDs, maximum and minimum wavelengths = 670 and 650, respectively; except for Experiment 1, see below) that bumblebees do not see well (Briscoe & Chittka 2001). Each colony contained ~50 worker bees and a queen at the beginning of each experiment. In order to obtain bees of known age, we collected callow (newly emerged, 0—24 hours of age) worker bees and marked each with a dot of acrylic paint-die (DecoArt, Stanford, KY, U.S.A, in Experiments 2 and 3) or a number-tag (Experiments 1, 4 and 5) on the dorsal part of their thorax. To obtain enough callow bees, collection, marking and reintroduction were done over two consecutive days (In Experiment 1, marking was done over 3 days). We used ‘white’ LED flesh lights (Energizer, St. Louis, Missouri, U.S.A) to facilitate callow-bee collection during day time (between 8:00 and 17:00). We re-collected the marked bees from their colonies (Experiments 1, 4 and 5) or experimental cages (Experiments 2, 3) when they were 5—7 days of age, and placed each individually in a monitoring cage. The individual cages were made of modified Petri dishes (diameter: 90 mm, height: 30mm) besides Experiment 1, in which we used different cages (see below). Each cage was provisioned with ad libitum pollen and sucrose syrup. After introducing into each cage the appropriate experimental stimulus (see below), we placed all the cages in a light-proof box and transferred them to an environmental chamber and recorded their locomotor activity (or behavior in Experiment 1).

### Locomotor activity monitoring and sleep analyses

We monitored the locomotor activity of individually isolated bees as previously described (Shemesh et al., 2007; Yerushalmi et al., 2006). Briefly, the monitoring cages with the focal bees and the experimental stimuli were placed in an environmental chamber (26—28°C, relative humidity 50%—70%) kept under constant dim red light (Edison Federal EFEF 1AE1 Far (Cherry) Red LED; maximum and minimum wavelengths were 750 and 730, respectively). Activity was recorded continuously over five successive days at a frequency of 1 Hz with four CCD cameras (Panasonic WV-BP334) and an image acquisition board (IMAQ 1409, National Instruments, U.S.A) laboratory. We estimated the proportion of daily sleep of individual bees, as described in Eban-Rothschild and Bloch (2015), on four consecutive days, starting on the 2^nd^ day of the monitoring session. Sleep was defined as a bout of five minutes or more in which the bee did not move. This sleep index was determined based on the detailed video recording analyses in Experiment 1 (see Results), and is consistent with similar sleep indices for honeybees (Eban-Rothschild and Bloch, 2008) and *Drosophila* (Shaw et al. 2000). We used the ClockLab circadian analyses software (Actimetrics, IL, USA) to generate *χ*^2^ periodograms using 10-minute bins and free running periods ranging between 20—28 hours. We used the ‘Power’ obtained from the periodogram as a proxy for the strength of circadian rhythmicity (Yerushalmi et al., 2006). The power was calculated as the height of the periodogram peak above the *p = 0.01* significance threshold. Bees with periodograms below the threshold line were assigned with a zero power value.

### Experiment 1: Detailed analyses of the behavior of workers isolated with or without pupae

On days 1—3 of the experiment, we marked focal callow bees from a single source colony. On Day 4, we changed the illumination regime to 12 hours light: 12 hours dark (LD; lights on at 07:00). During the light phase, the room was illuminated with standard florescent light (500 lux) and during the dark phase, with dim red light (as above). On days 7, 8, and 9, of the experiment, we transferred focal bees (4 bees in each day, marked on days 1, 2, and 3, respectively), into individual cages, such that all 12 bees were 6—7 days of age when isolated. The cages (7.5 × 5 × 3 cm) had a glass wall that enabled video recording. Each set of four individual cages was placed in a nearby environmental chamber under DD illumination regime. Bees were not exposed to light during the transfer. Two of the four cages had no brood (‘Brood-’). In each of these cages we placed a small piece of bumblebee wax obtained from a nectar pot in the source colony. Each of the remaining two cages received a single live intact pupa, also obtained from the same source colony (termed ‘Pupa+’). The developmental stage of these pupae was estimated based on inspections of at least three neighboring pupae and ranged from pre-pupa to purple-eyed pupa. On the following day after isolation of the focal bees, we video-recorded each of the four cages on four 1-hour sessions, using an infra-red video camera (Sony DCR-TRV75E). Two recordings were done during the subjective day (11:00 and 17:00; Group 1 was not recorded at 11:00) and two during the subjective night (23:00 and 5:00). We used the BORIS behavior analysis software (Friard & Gamba 2016) for detailed behavioral classification and time measurements. Table 1 summarizes the behaviors that we recorded in these analyses. One ‘Pupa+’ cage (from the 1 ^st^ set of cages) was excluded from the analysis because the introduced pupa was not properly attached to the substrate, turned loose, and disturbed the normal behavior of the focal bee. One broodless (‘Brood-’) bee from the same set was excluded because she displayed abnormal behavior with frequent atypical twitching and shivering. We additionally excluded the 5:00 observation of one ‘Pupa+’ bee in Group 3, because she was out of clear sight during most of the recording. At the end of video recordings, we kept the pupae in an incubator (33°, RH = 60%) until adult emergence, to confirm that the pupae were alive during the monitoring session. For a subset of ‘Sleep-like’ and ‘Incubation-like’ behavior bouts (see Table 1), in which we could clearly see the two antennae, we further recorded the movement of each antenna. We compared the frequency of antennal movements in bouts of these two behaviors from all bees using two-sample t-test. We also compared the bout length distributions for the three behaviors in which the bees were stationary: ‘Immobile-active’, ‘Sleep-like’ and ‘Incubation-like’ using the Kolmogorov-Smirnov test. Given that the distributions of bout length differed between the behaviors (see Results; Fig. 1C, D), we compared the bout lengths between the behaviors using the non-parametric Kruskal-Wallis test. We used Mann-Whitney tests with Bonferroni corrections for the *p-value* as *Post-Hoc* tests. In complementary χ^2^ tests of independence we limited our comparison to short (< 5 minutes) and long (≥ 5 minutes) bouts of immobility. We compared the proportion of recording sessions in which sleep was observed between bees with or without a pupa, using χ^2^ test of independence. The proportion of time spent sleeping and the number of sleep bouts per recording session were compared using two-sample t-tests. Given that sleep bout duration was not normally distributed, we compared bees with or without a pupa using the non-parametric Mann-Whitney test. We compared the proportion of time spent performing each behavior between the subjective day and subjective night with paired t-tests.

### Experiment 2: The effect of larvae on the sleep of tending workers

Fig. 2A describes the experimental design of Experiment 2. We paint marked all worker bees in 6 incipient colonies, and eight days later established 20 orphan (“queenless”) and 22 queenright cages. Into each queenless cage, we introduced two marked workers (> 8 days of age); into each queenright cage, we introduced a marked worker and a mated queen (obtained from Yad-Mordechai Pollination Services, Yad-Mordechai, Israel). We placed the bees in wooden cages with two removable glass walls (15 × 10 × 5 cm) and provisioned them with ad libitum pollen and sugar syrup. The cages were inspected daily in order to record new eggcups and monitor brood development. Six days later, when we begun the experiment, all the cages had larvae. On the next two days (Day 1 and 2; Fig. 2A), we introduced two additional paint marked sister callow-bees (the focal bees) into each one of the queenright and queenless cages. On Day 6, we collected the focal sister bees and placed each one individually in a monitoring cage with either a piece of wax (‘Brood-’) or ~8 live larvae sampled from her original cage (‘Larvae+’). Larvae from the queenright and the queenless cages were assumed to be females and males, respectively. This experimental design produced four experimental groups differing in the type of stimulus and the social environment experienced before isolation (Fig. 2A). The cages with the bees were then transferred to the locomotor activity monitoring-chamber. At the end of the monitoring session, we inspected the larvae from ‘Larvae+’ treatment and confirmed that they were alive. We compared the proportion of daily sleep between bees that were reared with or without larvae (Larvae factor) and bees that before monitoring experienced queenright or queenless conditions (Social environment factor, confounded with the larvae sex), using two-way ANOVA (SPSS) followed by LSD *Post-Hoc* tests.

### Experiment 3: The effects of larvae and pupae on the sleep of tending workers

We established 22 queenless cages, following the experimental setup described for Experiment 2. Twelve days later, when larvae developed in the queenless cages (Day 1 of the experiment), we introduced into each cage three marked sister callow workers (the “focal bees”) that were collected from 7 source colonies (see Fig. 2B). On Day 5, we collected pupae from the same source colonies and introduced one pupa into each queenless cage. On the next day (Day 6), each focal bee was collected from its queenless cage and placed in an individual monitoring cage together with either a piece of wax (‘Brood-’), ~8 larvae (‘Larvae+’) or a pupa (‘Pupa+’) that were collected from their original queenless cage (Fig. 2B). We then monitored the locomotor activity of the isolated focal bees, as described above. At the end of the monitoring session, we confirmed that the pupae and larvae were alive, as described for Experiments 1 and 2, respectively. Cages in which the pupa did not eclose were excluded from the analyses. We used one-way ANOVA followed by LSD *Post-Hoc* tests (SPSS) to assess the influence of treatment on the proportion of daily sleep.

### Experiment 4: The effects of live pupae and empty cocoons on the sleep of tending workers

The experimental setup is summarized in Figure 3A. Focal callow bees were marked and reintroduced to their source colonies on Days 1 and 2 of the experiment. On Day 7, we collected the focal bees from their source colonies, isolated each one of them in a separate monitoring cage and subjected them to one of 4 treatments. The ‘Brood-’ and ‘Pupa+’ treatments were similar to those described in Experiments 1, 2 and 3. In an additional treatment (termed ‘Empty’), we introduced an empty cocoon from which we gently removed the pupa through a longitudinal incision in the cocoon casing. In the sham control for this manipulation (‘Sham’), we made a similar incision in the pupal cocoon, but left the pupa intact. We used melted bumblebee wax to seal the incision in the cocoons of the ‘Empty’ and ‘Sham’ treatments. The focal bees in each of the three trials of this experiment were collected from 5—7 different source colonies and were assigned randomly to treatments. We monitored the locomotor activity of the bees over 5 days and then, using red light, entered the monitoring chamber and gently removed the treatment stimulus (i.e., ‘Brood-’, ‘Empty’, ‘Sham’, or ‘Pupa+’) from each one of the cages. We continued to monitor locomotor activity for additional five days (Fig. 3A bottom). The pupa containing cocoons were placed in an incubator (as above) until the emergence of callow bees to confirm vitality. Cages from which bees did not emerge were excluded from analyses. In Trials 2 and 3, we also recorded whether the focal bee started to construct wax pots. We used *χ*^2^ test of independence to test the effect of treatment on wax pot building. We used two-sample t-tests to compare the proportion of daily sleep time between bees that did or did not build wax pots using the pooled sample of all treatments, in Trials 2 and 3. The proportion of daily sleep was compared between treatments and between the days in the presence of treatments, using two-way repeated measures ANOVA followed by LSD *Post-Hoc* tests for the treatment factor. We also compared sleep proportion on day the 1^st^ and 4^th^ days of the monitoring session between treatments using one-way ANOVA and LSD *Post-Hoc* tests. Finally, we used paired t-tests to compare within each treatment, the proportion of sleep on the days before and after we removed the treatment stimuli.

### Experiment 5: The effect of pupal extracts on the sleep of tending workers

The basic design of this experiment is similar to Experiment 4 (Fig. S2A); we repeated this experiment three times. To prepare pupal extract, we collected fifteen cocoons with brood at prepupa to purple-eyed pupa stage from source colonies, and removed the top of each cocoon using fine scissors. We collected the cocoons and pupae and placed them into two separate glass Erlenmeyer flasks. We added 50 ml of n-Pentane (Sigma Aldrich, cat. #158941) into each Erlenmeyer, incubated the flasks at room temperature for 30 minutes, and poured the extract solution into a fresh 1000 ml Erlenmeyer flask. We washed each of the two extraction Erlenmeyer flasks again with 20 ml *n*-Pentane and added the washes into the 1000 ml Erlenmeyer flask. We repeated this procedure 4 times to obtain a total extract of 60 pupae and empty cocoons. We then suspended the 60 empty cocoons in 100 ml n-Pentane for additional 30 minutes at room temperature and added the extract to the stock solution in the 1000 ml Erlenmeyer flask. Finally, we similarly washed the empty cocoons one more time with n-Pentane in order to remove organic compounds. We concentrated the stock extract solution in a chemical hood for approximately 2 hours until its volume decreased to 30 ml, which constituted a concentration of two pupal equivalents/ ml. The washed cocoons were left to dry inside a chemical hood for ~24 hours. We then divided the stock solution with a glass pipette into twenty-eight 1-ml aliquots that were kept in glass vials, while alternately vortexing the solution to keep it well mixed. The aliquots were vortexed again and stored together with twenty eight 1ml pure *n*-Pentane aliquots at -20 °C until the next day.

On the following day, we thawed the extract aliquots, vortexed, and centrifuged them for 20 minutes at 4400 rpm. In Trial 1, we then applied the content of each extract solution or pure *n*-Pentane vial onto the surface of a washed cocoon, creating the ‘Extract’ and ‘Vehicle’ treatments, respectively. In Trials 2 and 3, we applied the vial content onto a 4 × 1 cm piece of filter paper (Trial 2) or a piece of cotton ball (Trial 3), in addition to the washed cocoon. The cocoons + filter paper / cotton-ball were then left in a chemical hood for ~2 hours until the solvent was evaporated. We then rolled and inserted the filter paper or cotton ball into the cocoon, which was then sealed with melted commercial sterilized honeybee wax (Nir Galim, Israel). These modifications in the protocols were done in an attempt to increase the bee exposure to the pupal extract. Each extract of vehicle lure was then placed in a monitoring cage together with a focal worker bee. As positive and negative controls, we used the ‘Brood-’ and ‘Pupa+’ treatments, as described above. We monitored locomotor activity during 5 consecutive days, as in Experiments 2, 3 and 4. At the end of the monitoring session, we opened the cocoons of pupae from the ‘Pupa+’ treatment and verified they were alive. As in Experiment 4, we compared the proportion of wax-pot building workers between treatments. We further used the χ^2^ test to compare wax pot building in the pooled sample of the negative control treatments (‘Brood-’ and ‘Vehicle’) compared to the ‘Pupa+’ treatment or the ‘Extract+’ treatment. We used a correction for the significance threshold (α) following (Sokal & Rohlf 1995): *α’* = 1 – (1 – *α*)^1/*k*^, where k = 2 is the number of comparisons. We tested the effect of wax building on daily sleep and the effects of Treatment and Day on sleep, as In Experiment 4.

### The relationship between ovarian development and sleep

A the end of the monitoring sessions, we dissected the abdomens of a random subset of focal bees (Experiment 3) or all focal bees (Trials 2 and 3 of Experiment 4) and measured the length of the largest ovarian oocyte, which was used as an index for ovarian state. We used two-way mixed-model ANOVA to test the effects of Live brood (pupae or larvae compared to the wax control; fixed factor), Experiment (random factor) and their interaction, on oocyte length. We excluded the empty cocoon treatment from this analysis. We next used multiple linear regression to test the effects of the experiment and the ovarian state on the proportion of daily sleep for all the treatments.

## Acknowledgements

We thank Inbal Tziperman-Lotan for her help in conducting the experiments, Dr. Hagai Shpigler for his assistance in the experimental design and Mira Cohen for technical support in the laboratory work. This work was supported by the Israel Science Foundation (ISF, number 1274/15 to GB).

