## Supplemental figures 1-3 for "Signals from the brood modulate the sleep of brood tending bumblebee workers"

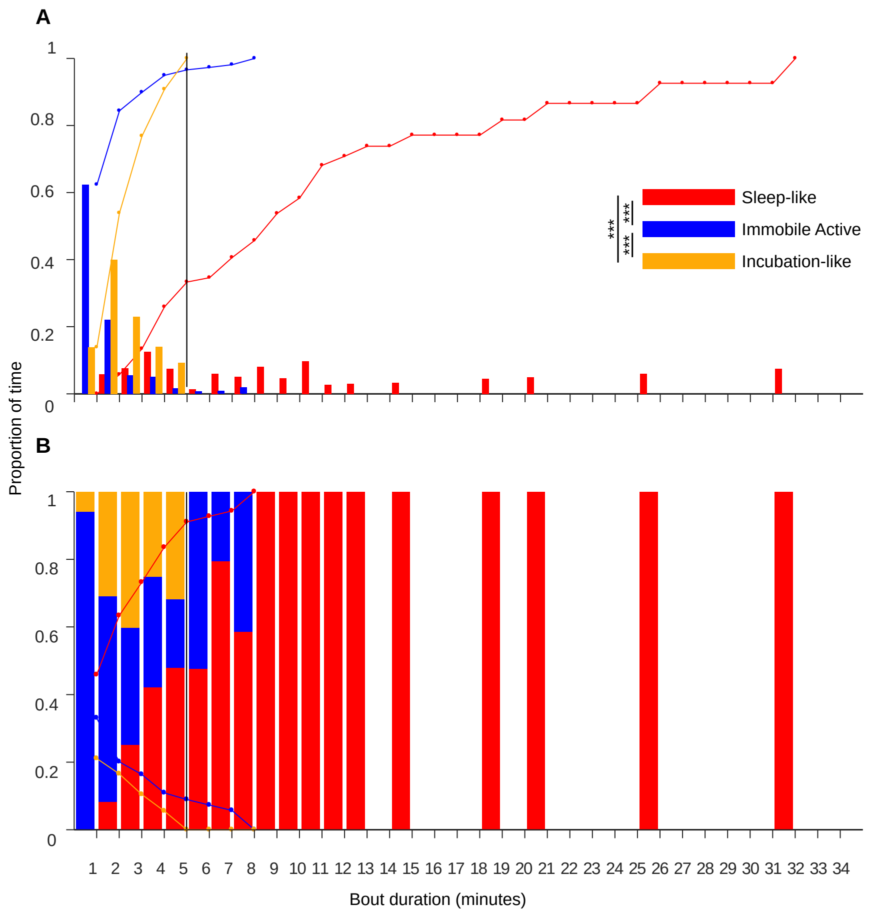


**Figure S1. Bout durations for immobile bees showing different behaviors.** The color legend marking the different behaviors is shown in panel A, and their description is provided in the text. (A) The frequency distribution (proportion of time) of bout durations in the different behaviors shown by immobile bees. The lines show the cumulative proportion of time. . The asterisks summarized the results of two-sample Kolmogorov-Smirnov test (***- *p* < *0.001*). (B) The relative proportion of time, in which immobile bees showed each behavior for a given bout duration. The lines show the accumulative proportions (≥). The vertical lines (A and B) indicate the 5' of immobility bout length threshold, which we used as a proxy for sleep bouts.

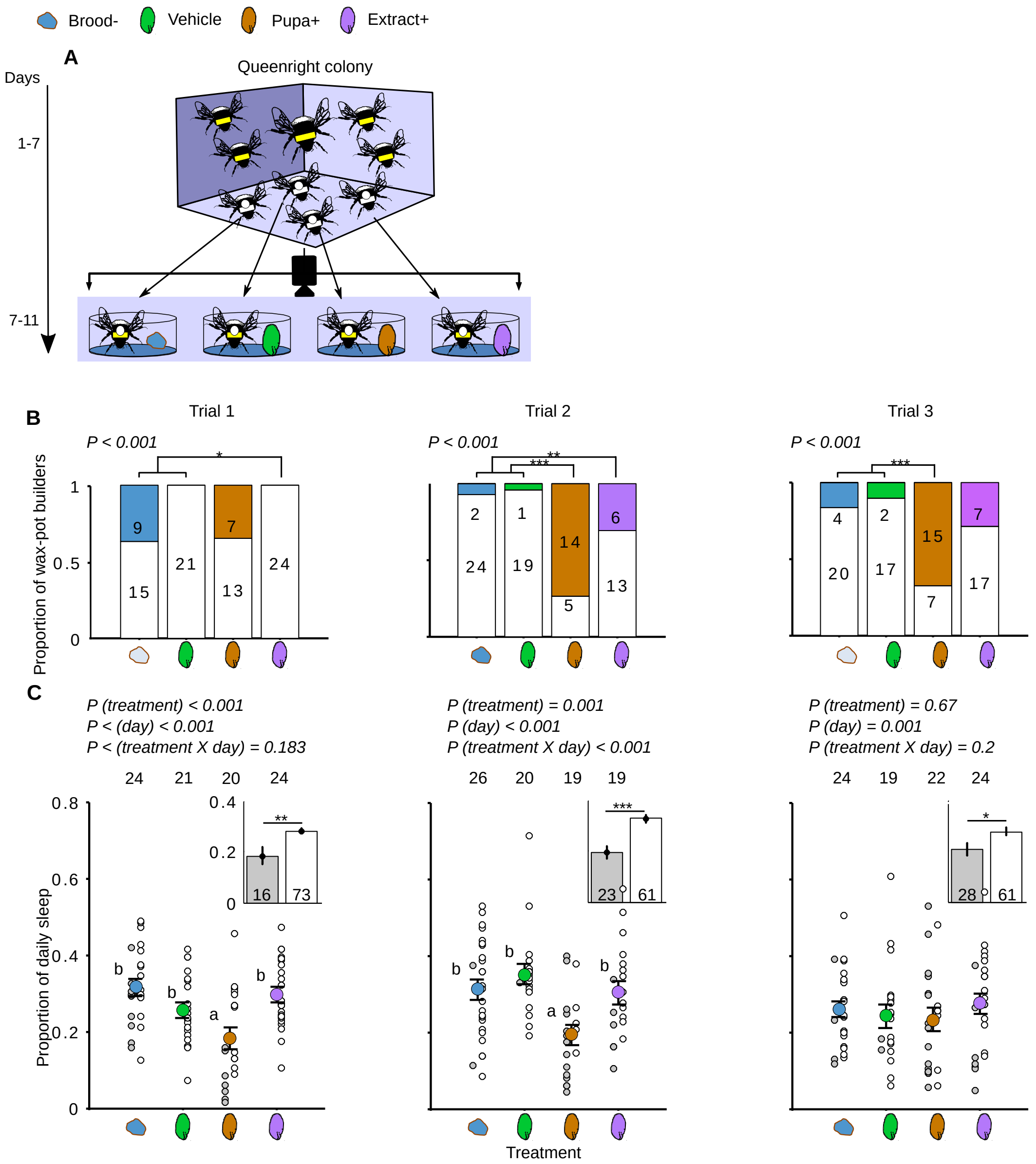


**Figure S2. The effect of cocoon extracts on sleep duration. (**A) The experimental setup of Experiment 5. Details of graph as in Fig. 3C, D with the following modifications: 'Extract+': a cocoon thoroughly washed with *n*-Pentane and treated with pupal extract; 'Vehicle' same, but treated with the pure *n*-Pentane solvent;. (B) The proportion of bees that initiated building wax pots. The *p*-values were obtained from 2 X 4 $\chi^2$ tests of independence followed by 2 X 2 complementary tests between the 'Pupa+' or 'Extract+' treatments and the pooled sample of the 'Vehicle' and 'Brood-' treatments. (C) The proportion of daily sleep. Other details as in Fig. 3.

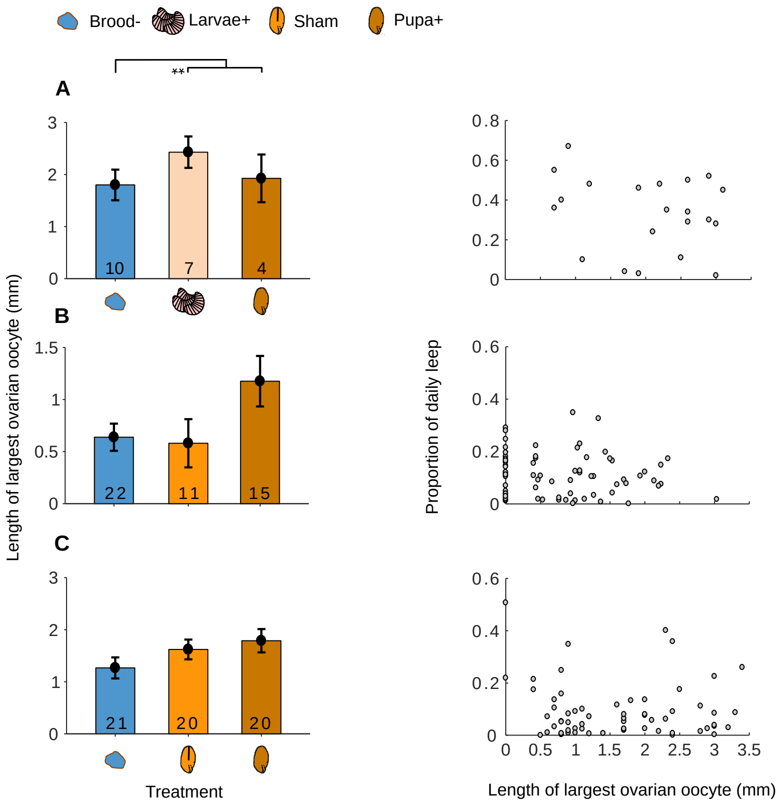


**Figure S3. The relationships between ovary state and sleep duration.** (A) Experiment 3; (B) Experiment 4, Trial 1; (C) Experiment 4, Trial 2. Left column: the length of the largest oocyte (means ± s.e.m, n) measured at the end of the experiment for bees in the presence of different treatment stimuli (details as in Fig. 2). Two-way ANOVA, $p > 0.001$. Right column: The relations between ovarian state (length of largest oocyte) and the proportion daily sleep.

**Movie 1: A bumblebee worker performing Incubation-like behavior.** The bee lays on-top of the pupa with her abdomen ‘extended’ over it, showing frequent pumping movements; her legs are wrapped around the pupa. The antennae are raised with an angle of > 90° between frons and scape, constantly moving without touching the substrate.

**Movie 2: A bumblebee worker performing sleep-like behavior.** The bee stops walking, gradually her antennae relax showing only low frequency movements, their angle falls below 90° between frons and scape**.** The ventilation pumping of the abdomen are discontinuous.
